# Double burden of malnutrition in children aged 24-59 months by socioeconomic status in five South Asian countries: evidence from Demographic and Health Surveys

**DOI:** 10.1101/605402

**Authors:** Fariha Binte Hossain, Md Shajedur Rahman Shawon, Md Shehab Uddin Al-Abid, Sultan Mahmood Sami, Gourab Adhikary, Md M Islam Bulbul

**Affiliations:** Independent Researcher; Nuffield Department of Population Health, University of Oxford, Richard Doll Building, OX3 7LF, UK; National Heart Foundation Hospital and Research Institute, Dhaka, Bangladesh; Health Systems and Population Studies Division, icddr,b, 68 Shaheed Tajuddin Ahmed Sarani, Mohakhali, Dhaka 1212, Bangladesh; National Nutrition Services, Ministry of Health and Family Welfare, Bangladesh

**Keywords:** Double burden, underweight, overweight, under-five children, South Asia

## Abstract

**Background:** Developing countries are now facing double burden of undernutrition and overnutrition among children and adults. We aimed to explore the double burden of malnutrition among children aged 24-59 months by household’s socioeconomic status in South Asian context.

**Methods:** Children with valid information on height and weight from the latest Demographic and Health Survey from Bangladesh, India, Pakistan, Maldives, and Nepal were included in this study. Underweight and overweight were defined according to definitions of World Health Organisation and International Obesity Task Force, respectively. We used multiple logistic regressions to estimate the association of socioeconomic status with childhood underweight and overweight.

**Results:** South Asian countries had significant burden of underweight, ranging from 19% in Maldives to 38% in India. Bangladesh, India, and Nepal had prevalence of overweight between 2% and 4%, whereas Pakistan and Maldives had prevalence of 7% and 9%, respectively. Households with higher wealth index and education were consistently associated with lower odds of underweight children. When compared to poorest households, richest households had higher odds of being overweight in Bangladesh (OR 1.96, 95% CI 1.27-3.02) and India (OR 1.53, 95% CI 1.41-1.66) while lower odds of being overweight in Pakistan (OR 0.22, 95% CI 0.14-0.34). Households with higher education were more likely to have overweight children in Bangladesh and India.

**Conclusions:** Childhood underweight is associated with lower socioeconomic conditions while there is a substantial burden of childhood overweight in higher socioeconomic groups. These disparities by socioeconomic conditions should be considered while developing national nutrition programs and strategies.

**KEY MESSAGES:**

- In South Asia, there is a substantial burden of undernutrition among under-five children while a differential burden of overnutrition is also seen.
- Household wealth and educational attainment were inversely associated with childhood underweight.
- Children in households with higher levels of wealth and educational attainment were more likely to be overweight in Bangladesh and India, while evidence supporting such association was not clear for other South Asian countries.
- The urban-rural difference in the burden of childhood underweight and overweight can be explained by the distributions of households’ socioeconomic status.

## INTRODUCTION

Double burden of malnutrition implies to the presence of both undernutrition and overnutrition (overweight or obesity) within individuals, households, or populations.^1^ At the individual level an undernourished child can be overweight or obese when they reach adulthood, whereas at household level coexistence of underweight and overweight children or adults can be possible. At the population level, double burden of malnutrition indicates the prevalence of both underweight and overweight in the same community, country, or region.^1,2^

In low and middle-income countries (LMICs), underweight in children under the age of five years has been a major public health problem historically.^3,4^ South Asia is comprised mostly of LMICs, and had highest numbers of underweight children due to higher prevalence rates and large populations in younger age groups.^5^ However, due to recent economic growth, rapid urbanization, and adoption of western lifestyles, the burden of overweight in this age group is increasing in LMICs, including South Asian countries.^6–8^ Also, South Asians children living in developed countries have a higher prevalence of overweight than any other ethnic group - a recent study suggests.^9^ So far health systems in South Asia are focusing mainly on prevention of childhood undernutrition, but an increasing trend in overnutrition demands newer approach to tackle double burden of malnutrition among children in this region. Both childhood underweight and overweight are associated with wide range of morbidities in early life as well as in later life.^6,10^

To have better strategies to solve the problem of double burden of disease among children under the age of five years, we need to understand the socioeconomic inequalities in nutritional outcomes. Identifying socioeconomic groups with higher burden of nutritional problems can help tailoring public health prevention interventions. Previous studies have suggested that there can be substantial differences in the burden of malnutrition by household’s wealth, education level, and area of residence.^11–14^ Although the double burden of malnutrition has received much attention in recent time, there is no study, to the best of our knowledge, which looked at this problem in relation to various measures of socioeconomic status in nationally-representative samples.

In this study, we aimed to quantify the extent of underweight and overweight among children aged 24-59 months in South Asian countries and to estimate their associations with household’s socioeconomic status, using the latest nationally-representative surveys.

## METHODS

### Study design and data sources

This study is based on the latest DHS data from five South Asian countries, namely Bangladesh, India, Pakistan, Maldives, and Nepal. Other countries in the South Asian regions (e.g. Afghanistan, Bhutan, and Sri Lanka) were not included in this study because of either DHS was not conducted, or anthropometric data for children were not available.

DHS are nationally-representative household surveys which are usually conducted about every 5 years. These surveys provide data for a wide range of monitoring and impact evaluation indicators in the areas of population, health, and nutrition. A DHS is conducted by a national implementing agency, which can be any bona-fide governmental, non-governmental, or private-sector organization and has enough experience in the execution of surveys that are national in scope. Technical assistances throughout the whole process are provided by the DHS program.^15^ DHS are usually based on two-stage stratified sampling of households. In the first stage, sampling census enumeration areas are selected using probability proportional to size (PPS) sampling technique through statistics provided by the respective national statistical office. In the second stage, households are selected through systematic random sampling from the complete listing of households within a selected enumeration area.^16^

Ethical approval for each DHS is taken from the ICF Institutional Review Board as well as by a review board in the host country. More details of such ethical approval can be found in the DHS program website [https://dhsprogram.com/]. Informed consent to participate in the study is taken from the participant, or from the parent or guardian if anthropometric measurements are taken from a child. The data files are freely available from the program website. We received authorization from the DHS program for using the relevant datasets for this analysis. The data we received were anonymized for protection of privacy, anonymity and confidentiality.

These surveys have very high response rate, usually 90% and above. Detailed questionnaires of included surveys are available in the final report of each survey. We used the children’s record (coded as “KR” in DHS program) datasets which contained information about children born in the last 5 years prior to the survey (aged 0-59 months). The present analysis is based on children aged 24 – 59 months who had valid measurement of their weight and height. We excluded children aged less than 24 months because there is no available classification system for defining overweight for them.

### Anthropometric measurement, and defining underweight and overweight

In DHS, height and weight of the children were measured by trained personnel using standardized instruments and procedures. Lightweight SECA scales (Hamburg, Germany) with digital screen, designed and manufactured by the United Nations Children’s Fund (UNICEF), were used to measure weight. The height/length was measured by boards, produced by Shorr Productions (Maryland, USA). In children with height less than 85 centimetres, recumbent length was measured, whereas standing height was measured for those taller than this. Body mass index (BMI) was calculated by dividing body weight (kg) by squared height (m^2^).

Childhood underweight is based on the indicator weight-for-age, which is an overall indicator of population’s nutritional status. A child with weight-for-age less than two standard deviations (−2 SD) from the median of the reference population is considered as underweight according to World Health Organization (WHO) guidelines.^17^ Underweight is a composite definition which can encompass stunting, wasting, or both.

To define childhood overweight, we used the age and sex-specific BMI cut-offs from the International Obesity Task Force (IOTF) classification system.^18,19^ According to IOTF, a child aged between 2 years and 18 years is classified as overweight if their BMI is larger than the age and sex-specific BMI cut-off corresponding to an adult BMI of >25 kg/m^2^. Our definition of childhood overweight also included those with obesity and it is referred to hereafter as “overweight” for simplicity.

### Socioeconomic factors

DHS collected information on wide range variables from the selected households and the respondents from those households using face-to-face interview conducted by trained personnel. DHS collected information on socioeconomic factors like area of residence and household’s wealth index. Place of residence (rural and urban) was defined according to country-specific definitions. For household’s wealth index, each national implementing agency constructed a country-specific index using principal components analysis from data on household assets including durable goods (i.e. bicycles, televisions etc.) and dwelling characteristics (i.e. sanitation, source of drinking water and construction material of house etc.).^15^ This wealth index was then categorized into five groups (i.e. poorest, poorer, middle, richer, and richest) based on the quintile distribution of the sample.

### Statistical analysis

We conducted all analysis following the instructions given in the DHS guide to analysis.^16^ The percent distributions for characteristics of included children are described as proportions, for each DHS survey. To estimate the prevalence of childhood underweight and overweight, we used sampling weights given in each DHS dataset in order to get nationally-representative estimates. 95% confidence intervals (CIs) for prevalence estimates were calculated using a logit transform of the estimate. We also estimated the prevalence of childhood underweight and overweight by the levels of socioeconomic factors to assess the inequalities by those factors.

To examine the associations of socioeconomic factors with prevalence of childhood underweight and overweight, we used multiple logistic regression, separately for each included country. These analyses were adjusted for child’s age, sex, are of residence, household’s highest education level, household’s wealth index, as appropriate. Considering the two-stage stratified cluster sampling in DHS, we applied Stata’s survey estimation procedures (*“svy”* command) for regression analyses.^20^ The results are presented (as in tables and figures) as group-specific 95% confidence intervals (g-SCIs) for comparisons between more than two categories to allow comparisons to be made between any two categories, even if neither is the reference group.^21^ Conventional 95% CIs are provided in case of two categories being compared. All analyses were performed using Stata v15.1 (Statacorp, College Station, TX, USA).

## RESULTS

A total of 146 996 children aged between 24 and 59 months from five south Asian countries were included in this study. Table 1 shows the sample characteristics for each of these countries’ latest DHS data. Sample population for five countries had almost equal distribution for both sex and age. In all countries except Nepal, the majority of the children were from rural area (according to the definition of specific country), and the proportions varied widely between 57% and 86%. On average less than one in every 10 households had at least one member who completed higher education. India, Nepal and Pakistan had significant number (≥33%) of households where none had formal education, whereas in Bangladesh and Maldives proportion of such households was relatively lower (<20%). The samples from original surveys were divided into quintiles based on household’s wealth index, and after relevant exclusions for this study the distributions remained more or less similar (Table 1). As expected, the better part of burden for malnutrition in all countries was due to undernutrition (Figure 1). India had the highest (38%) prevalence of undernutrition among children aged 24-59 months followed by Bangladesh (37%), Nepal (29%), Pakistan (28%), and Maldives had the lowest prevalence (19%). For overweight among these children, Bangladesh, India, and Nepal had similar prevalences (between 2% and 4%) whereas Pakistan and Maldives had much higher prevalence, 7% and 9% respectively. When we looked at the combined burden of both forms of malnutrition, India and Bangladesh had much higher burden in compared to other countries. However, the prevalences for both undernutrition and overnutrition were slightly higher in India than those in Bangladesh. In Pakistan and Maldives where overweight prevalence was high, the burden of undernutrition was decreased.

**Table 1:** Sample characteristics in five demographic and health survey data, by country.

|  | Number (%) |  |  |  |  |
| --- | --- | --- | --- | --- | --- |
|  | Bangladesh | India | Maldives | Nepal | Pakistan |
| Year of survey | 2014 | 2015-16 | 2009 | 2016 | 2012-13 |
| Number of children | 4170 | 138134 | 1339 | 1389 | 1964 |
| Child's sex |  |  |  |  |  |
| Male | 2134 (51.2) | 71698 (51.9) | 672 (50.2) | 715 (51.5) | 1016 (51.7) |
| Female | 2036 (48.8) | 66436 (48.1) | 667 (49.8) | 674 (48.5) | 948 (48.3) |
| Child's age |  |  |  |  |  |
| 2 year | 1406 (33.7) | 45298 (32.8) | 452 (33.8) | 460 (33.1) | 668 (34.0) |
| 3 year | 1377 (33.0) | 47506 (34.4) | 464 (34.7) | 479 (34.5) | 641 (32.6) |
| 4 year | 1387 (33.3) | 45329 (32.8) | 423 (31.6) | 449 (32.3) | 655 (33.4) |
| Area of residence |  |  |  |  |  |
| Urban | 1316 (31.6) | 33245 (24.1) | 183 (13.7) | 788 (56.7) | 851 (43.3) |
| Rural | 2854 (68.4) | 104889 (75.9) | 1156 (86.3) | 601 (43.3) | 1113 (56.7) |
| Household's highest education level |  |  |  |  |  |
| No education | 714 (17.1) | 44950 (32.5) | 221 (16.5) | 514 (37.0) | 1067 (54.3) |
| Primary | 1168 (28.0) | 20664 (15.0) | 615 (45.9) | 260 (18.7) | 303 (15.4) |
| Secondary | 1877 (45.0) | 60737 (44.0) | 462 (34.5) | 431 (31.0) | 385 (19.6) |
| Higher | 411 (9.9) | 11783 (8.5) | 26 (1.9) | 184 (13.2) | 209 (10.6) |
| Wealth index |  |  |  |  |  |
| Poorest | 931 (22.3) | 36404 (26.4) | 330 (24.6) | 351 (25.3) | 443 (22.6) |
| Poorer | 781 (18.7) | 32673 (23.7) | 335 (25.0) | 308 (22.2) | 390 (19.9) |
| Middle | 808 (19.4) | 27462 (19.9) | 358 (26.7) | 296 (21.3) | 323 (16.4) |
| Richer | 843 (20.2) | 23044 (16.7) | 201 (15.0) | 276 (19.9) | 419 (21.3) |
| Richest | 807 (19.4) | 18551 (13.4) | 115 (8.6) | 158 (11.4) | 389 (19.8) |

**Figure 1:**
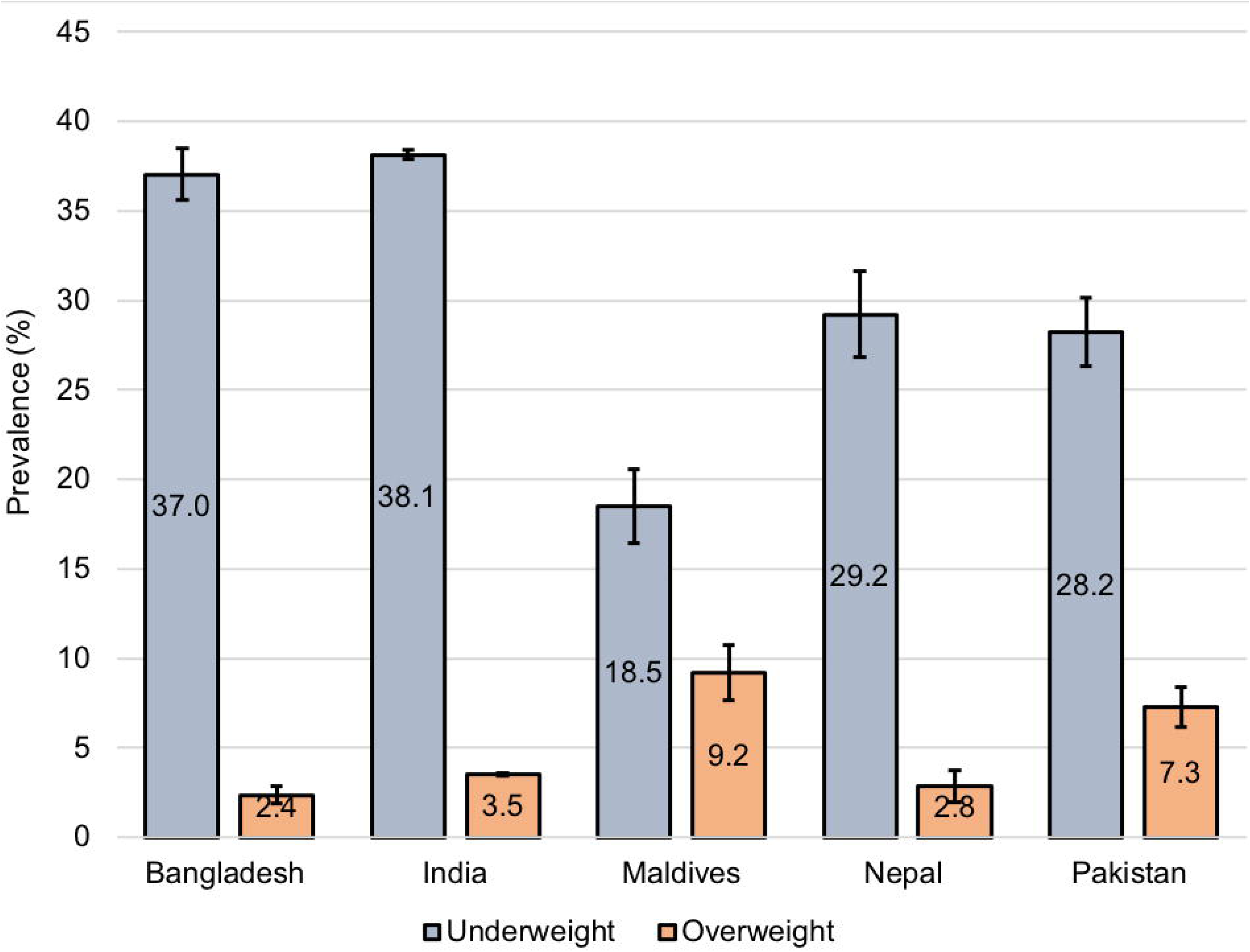
Prevalence of underweight and overweight, by country Sampling weight provided by the Demographic and Health Survey (DHS)^16^ was used to estimate country-representative prevalence. Error bars represent 95% confidence intervals

There were wide variations in the prevalence of undernutrition and overweight according to household’s wealth index (Figure 2) and household’s highest education (Figure 3) in all countries. Between the poorest and the richest households, the burden of undernutrition decreased by more than half in all of these five countries. The prevalence of overweight almost doubled between the poorest and the richest households in Bangladesh and India, whereas such differences were not clearly evident in Maldives and Nepal. In Pakistan, rich households were less likely to have overweight children than poor households. The prevalence of undernutrition and overweight according to household’s highest education level followed similar country-specific patterns observed for wealth index (Figure 3). Notably, children in households with higher education had much higher rate of overweight in Bangladesh, India, and Pakistan.

**Figure 2:**
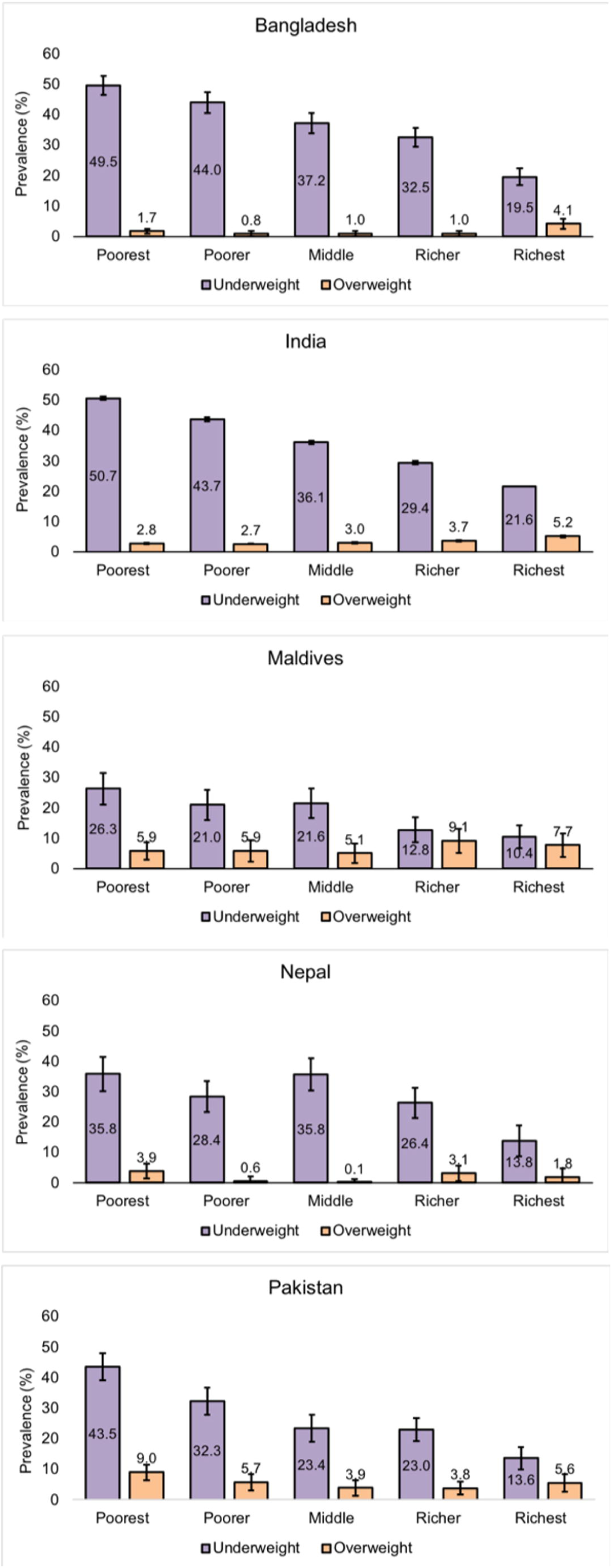
Prevalence of underweight and overweight, by household’s wealth index Error bars represent 95% confidence intervals

**Figure 3:**
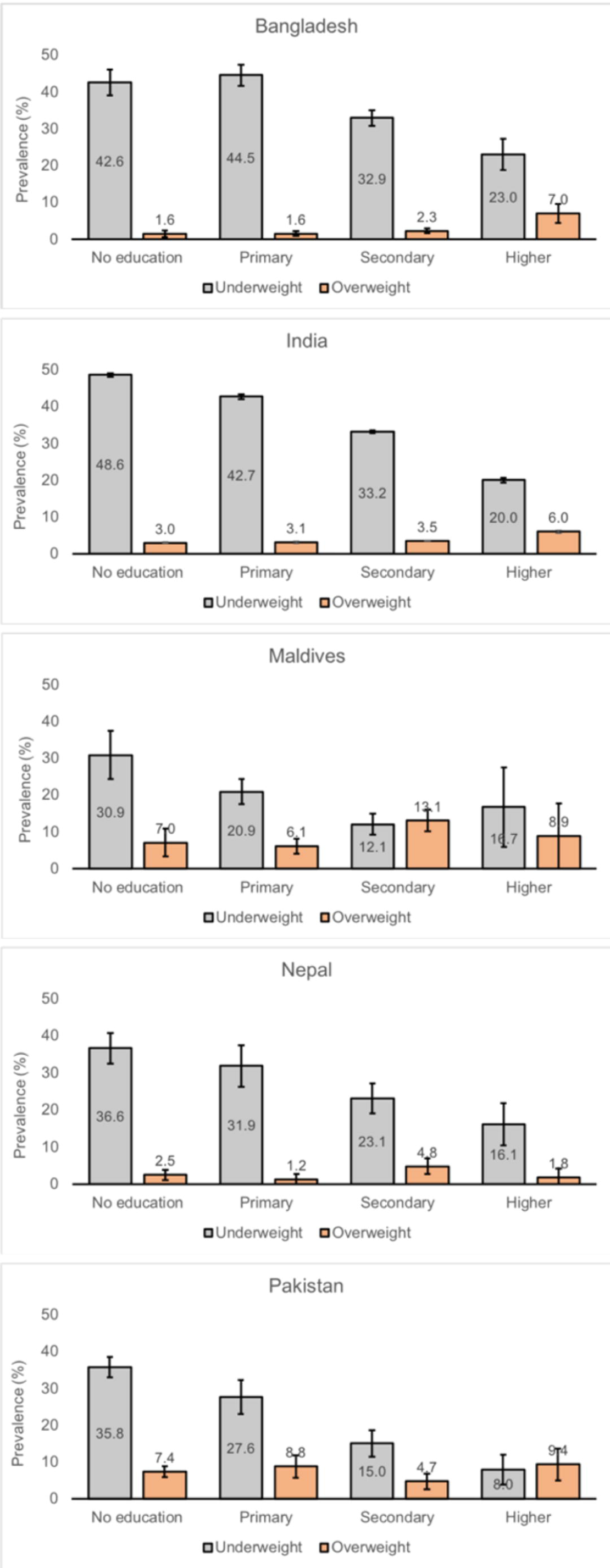
Prevalence of underweight and overweight, by household’s highest educational attainment Error bars represent 95% confidence intervals

When adjusted for age, sex, area of residence, and education, there was reliable evidence for inverse relationship between wealth index and prevalence of underweight in Bangladesh, India, Nepal, and Pakistan; whereas such conclusion could not be made for Maldives possibly due to smaller number of cases (Figure 4). The adjusted ORs for richest vs. poorest households were 0.36 (95% CI 0.34-0.37) and 0.31 (95% CI 0.25-0.37) in India and Bangladesh, respectively. Richest households were more likely to have children who were overweight in compared to the poorest households in India (OR 1.53, 95% CI 1.41-1.66) and in Bangladesh (OR 1.96, 95% CI 1.27-3.02). In contrary, richest households were less likely to have overweight children in Pakistan when compared to poorest households (OR 0.22, 95% CI 0.14-0.34). The overall associations of wealth index with underweight and overweight were not significant for Maldives and Nepal.

**Figure 4:**
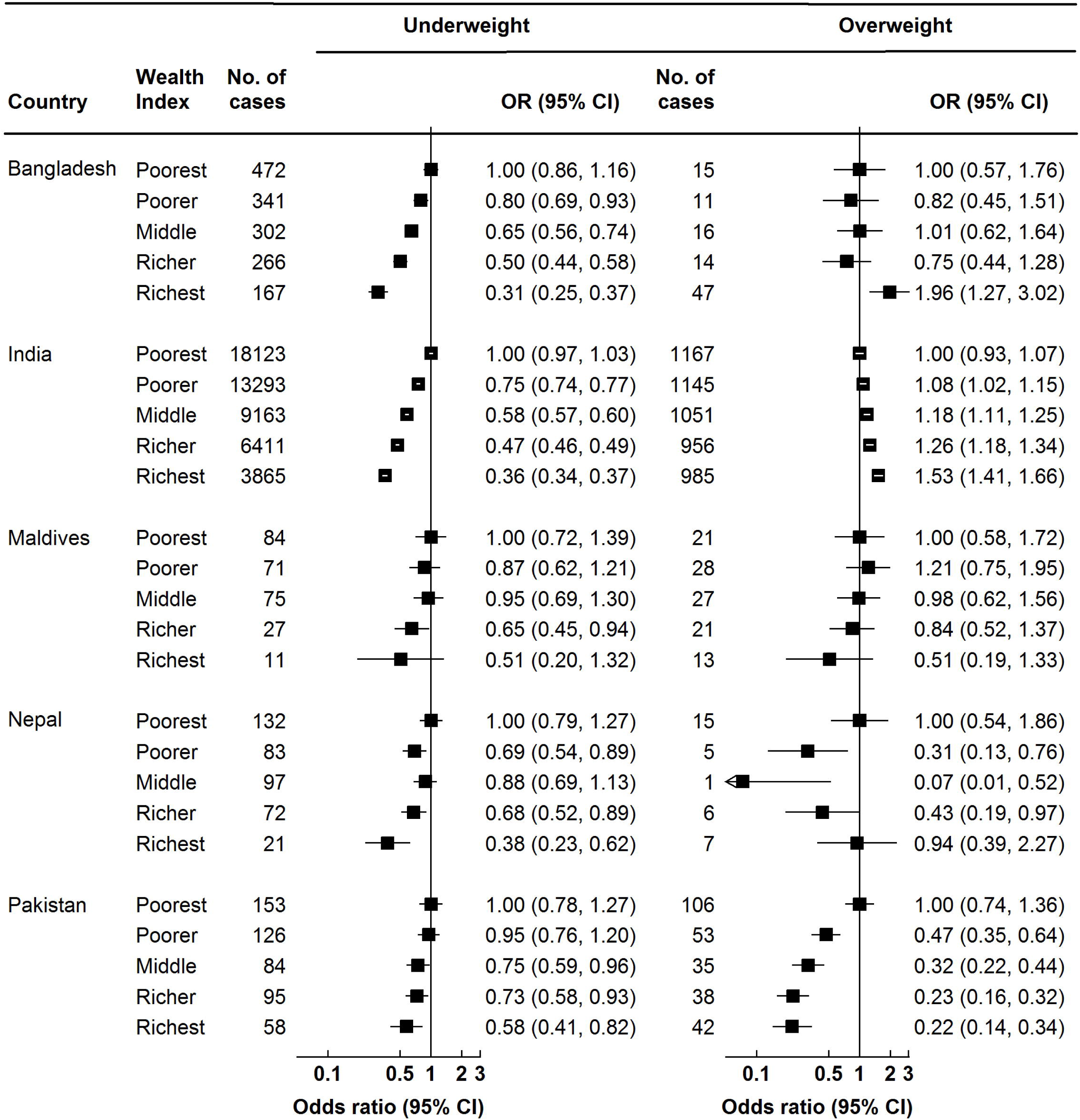
Odds ratios of underweight and overweight, by household’s wealth index Analyses were adjusted for age, sex, area of residence, and household’s highest educational attainment

There were significant inverse associations between household’s education level and underweight in all countries expect Maldives, after adjustment for age, sex, area of residence, and wealth index (Figure 5). The ORs of underweight for higher education vs. no education were 0.48 (95% CI 0.46-0.51), 0.63 (95% CI 0.49-0.81), and 0.38 (95% CI 0.24-0.59) in India, Bangladesh, and Pakistan, respectively. Households with higher education were more likely to have overweight children when compared to households with no education in Bangladesh (OR 2.97, 95% CI 1.88-4.68), and in India (OR 1.37, 95% 1.25-1.50). Overweight in children was not associated with education level in Maldives, Nepal, and Pakistan.

**Figure 5:**
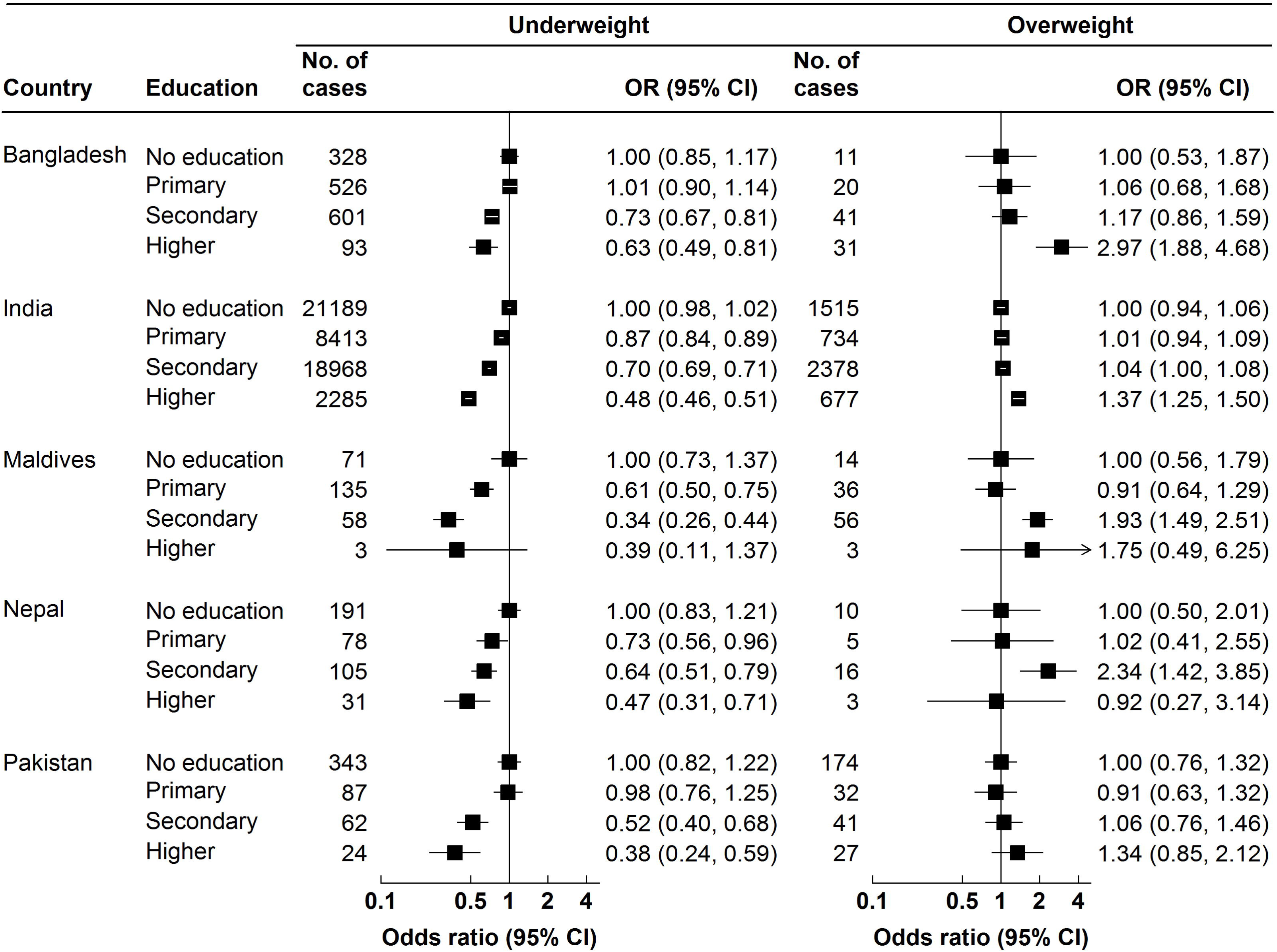
Odds ratios of underweight and overweight, by household’s highest educational attainment Analyses were adjusted for age, sex, area of residence, and household’s wealth index

Additional analyses showed that there were no appreciable sex differences for underweight and overweight prevalence (Supplementary Table S1). The prevalence for underweight and overweight differed between rural and urban areas (Supplementary Table 2), although adjusted models showed no significant association for area of residence with underweight and overweight (Supplementary Table 3). This illustrates that socioeconomic status can explain the rural-urban differences in double burden of malnutrition.

## DISCUSSION

This study involving nationally-representative surveys conducted in recent times in five South Asian countries provided empirical evidence on double burden of malnutrition among children aged 24-59 months and its association with socioeconomic status factors. In South Asian countries, there is a substantial burden of undernutrition among younger children while a differential burden of overnutrition is also seen. Households with higher socioeconomic status (as measured by wealth index and education) were consistently associated with lower odds of underweight children in all countries, though the association did not reach statistical significance in Maldives. The associations between socioeconomic status and overweight were heterogeneous: both households with richest wealth and households with higher education were more likely to have overweight children in India and Bangladesh, but the evidence for such associations in other countries was not consistent.

South Asian countries have experienced striking economic growth in the last few decades which triggered unprecedented improvements in maternal mortality, infant mortality, under-five mortality, and child undernutrition.^22,23^ The prevalence of childhood underweight was declined by 25-30% between 2004 and 2014 in Bangladesh, India, Pakistan, and Nepal.^24^ However, the existing burden of undernutrition is still high – our study found that around one-third of under-five children in this region are underweight. Previous studies conducted in the region have found that poor socioeconomic status, lower level of parental education, younger age of mother at birth, short birth interval, and initiation of complimentary feeding are important determinants of undernutrition among under-five children.^25–27^ We also observed significant nutrition disparity by household socioeconomic status. In populous countries like Bangladesh, India, and Pakistan, almost half of the children in poorest households were underweight. In multivariable models, both household’s socioeconomic status and household’s highest education level were found to be strongly associated with childhood underweight in all countries. Multi-sector approach is needed to alleviate poverty and other social inequalities related to undernutrition disparity in South Asia and beyond.^28^

Recent reports^29–32^ from South Asian countries highlighted the rise of overweight burden in children, but mainly in older groups. Overweight among under-five children is still overlooked in current literature. In our study, we provided evidence for an increasing burden of overweight in this age group, which clustered in households with higher socioeconomic status. We used two indicator variables for household’s socioeconomic status, namely wealth index and highest level of education, and found that after simultaneous adjustment for each other wealth index had better explanatory power than education level. Frequent intake of energy-dense foods and physical inactivity have been shown to be associated with overweight and obesity both in children and adults.^33,34^ These lifestyle behaviors are common in the higher socioeconomic group in LMICs and therefore, both childhood and adulthood overweight are clustered in affluent households in urban areas.^29,32^ Public health nutrition programmes should therefore focus on educating parent of younger children about proper feeding guidance and importance of physical activity.

South Asian countries have seen an unprecedented rise of urbanization and economic growth in recent times.^35^ Previous studies^29,32,36^ reported about urban-rural gap in burden of overweight and obesity, but we found no significant differences after adjustment for socioeconomic variables. This means that socioeconomic distribution of households can explain the observed urban-rural differences for the burden of childhood overweight. In our study, the associations for socioeconomic status with childhood overweight were heterogeneous among countries, but it could be due to small number of overweight children in those countries. A previous study from Pakistan with a representative multistage cluster sample also found that affluent urban population was facing a rapid rise in overweight and obesity among primary school children.

The findings from our study highlight the importance of considering social determinants of health while developing public health interventions and policies to tackle both childhood undernutrition and overnutrition. So far the public health interventions in South Asia focus almost completely on the prevention of undernutrition, but identifying groups with more likelihood of developing childhood overweight and obesity can help to shift the focus of intervention to those groups. We suggest the policy makers to provide more resources to tackle underweight while care should be taken for the affluent section of the society to prevent overweight. To the best of our knowledge, no study looked at the coexistence of underweight and overweight among under-five children in South Asian countries by socioeconomic status. We used nationally-representative samples for five South Asian countries to investigate the association of double burden of malnutrition with households’ socioeconomic status. Child’s height and weight were measured objectively by trained field researchers using calibrated scales. We also used IOTF classification system to define overweight among these children, which helps to compare the overweight prevalence internationally. We were also able to adjust for several factors in the multivariable models. Our study lacks information on dietary and lifestyle factors, so we could not adjust for their effects on the association between socioeconomic status and double burden of malnutrition. Due to smaller sample sizes in Maldives and Nepal, we could not reliably estimate the associations. We excluded those children who did not have anthropometric data, but DHS reports suggest that they should not vary significantly in terms of sociodemographic characteristics.

In conclusion, our study provides evidence for socioeconomic disparities for the coexistence of under and over nutrition among children aged 24-59 months in South Asian settings. These unmet inequalities for both underweight and overweight should be considered while developing national public health nutrition programmes and strategies.

## ACKNOWLEDGEMENTS

The authors thank the participants of Demographic and Health Surveys used in this study from Bangladesh, India, Maldives, Nepal, and Pakistan. We would also like to thank the DHS Program to authorize us to use the data.

## AUTHOR CONTRIBUTIONS

Conception and design: FH, MS, SA and MB

Data collection and management: FH, SS, and GA

Data analysis: FH, MS, SS

Interpretation of the results: All authors

Drafting of the article: FH and MS

Critical revision of the article for important intellectual content: All authors

Final approval of the article: All authors.

