## Supplementary materials for "Double burden of malnutrition in children aged 24-59 months by socioeconomic status in five South Asian countries: evidence from Demographic and Health Surveys"

**Table S1: Prevalence of underweight and overweight among children aged 24-59 months, by sex**

|  | **Prevalence (95% CIs)** | |
| --- | --- | --- |
|  | **Boys** | **Girls** |
| **Underweight** |  |  |
| Bangladesh | 35.2 (33.2-37.2) | 39.0 (36.9-41.1) |
| India | 37.4 (37.0-37.7) | 38.9 (38.6-39.3) |
| Maldives | 17.6 (14.9-20.7) | 19.4 (16.6-22.6) |
| Nepal | 29.3 (26.1-32.8) | 29.1 (25.8-32.6) |
| Pakistan | 30.6 (27.9-33.4) | 25.7 (23.1-28.4) |
| **Overweight** |  |  |
| Bangladesh | 2.0 (1.5-2.6) | 2.8 (2.1-3.6) |
| India | 3.4 (3.2-3.5) | 3.7 (3.5-3.8) |
| Maldives | 8.3 (6.5-10.7) | 10.0 (7.9-12.5) |
| Nepal | 1.7 (1.0-3.0) | 4.0 (2.7-5.7) |
| Pakistan | 7.2 (5.8-8.9) | 7.3 (5.9-9.1) |

Sampling weights provided by the Demographic and Health Survey (DHS) were used to estimate country-representative prevalence. 95% confidence intervals (CIs) for prevalence estimates were calculated using a logit transform of the estimate.

**Table S2: Prevalence of underweight and overweight among children aged 24-59 months, by area of residence**

|  | **Prevalence (95% CIs)** | |
| --- | --- | --- |
|  | **Urban** | **Rural** |
| **Underweight** |  |  |
| Bangladesh | 30.0 (27.4-32.9) | 39.4 (37.7-41.1) |
| India | 30.9 (30.4-31.4) | 41.0 (40.7-41.3) |
| Maldives | 11.5 (8.8-15.0) | 21.7 (19.1-24.4) |
| Nepal | 25.4 (22.4-28.7) | 33.6 (30.1-37.4) |
| Pakistan | 23.3 (20.2-26.7) | 30.4 (28.1-32.8) |
| **Overweight** |  |  |
| Bangladesh | 3.9 (2.9-5.2) | 1.8 (1.4-2.4) |
| India | 4.4 (4.2-4.7) | 3.1 (3.0-3.2) |
| Maldives | 13.2 (10.2-16.8) | 7.4 (5.8-9.2) |
| Nepal | 3.1 (2.1-4.6) | 2.5 (1.5-4.0) |
| Pakistan | 6.6 (4.9-8.8) | 7.6 (6.3-9.1) |

Sampling weights provided by the Demographic and Health Survey (DHS) were used to estimate country-representative prevalence. 95% confidence intervals (CIs) for prevalence estimates were calculated using a logit transform of the estimate.

**Table S3: Associations of area of residence with underweight and overweight among children aged 24-59 months**

|  | **Area** | **No. of case** | **Adjusted OR (95% CIs) ^†^** |
| --- | --- | --- | --- |
| **Underweight** |  |  |  |
| Bangladesh | Urban | 416 | 1.00 (Reference) |
|  | Rural | 1132 | 0.94 (0.80-1.10) |
| India | Urban | 10075 | 1.00 (Reference) |
|  | Rural | 40780 | 0.90 (0.87-0.92) |
| Maldives | Urban | 20 | 1.00 (Reference) |
|  | Rural | 248 | 1.03 (0.46-2.31) |
| Nepal | Urban | 204 | 1.00 (Reference) |
|  | Rural | 201 | 1.11 (0.87-1.43) |
| Pakistan | Urban | 185 | 1.00 (Reference) |
|  | Rural | 331 | 1.01 (0.79-1.29) |
| **Overweight** |  |  |  |
| Bangladesh | Urban | 52 | 1.00 (Reference) |
|  | Rural | 51 | 0.69 (0.44-1.09) |
| India | Urban | 1420 | 1.00 (Reference) |
|  | Rural | 3884 | 1.05 (0.98-1.13) |
| Maldives | Urban | 25 | 1.00 (Reference) |
|  | Rural | 85 | 0.39 (0.16-0.96) |
| Nepal | Urban | 18 | 1.00 (Reference) |
|  | Rural | 16 | 1.23 (0.57-2.66) |
| Pakistan | Urban | 118 | 1.00 (Reference) |
|  | Rural | 156 | 0.57 (0.41-0.79) |

^†^ Adjusted for age, sex, household’s highest education and household’s wealth index
